# Sparse spatial scaffolding for visual working memory

**DOI:** 10.1101/2023.07.05.547765

**Authors:** Baiwei Liu, Zampeta-Sofia Alexopoulou, Siyang Kong, Anne Zonneveld, Freek van Ede

## Abstract

When holding information ‘in mind’, it is vital to keep individual representations separated and selectively accessible for behaviour. Space is known to serve as a foundational scaffold for mnemonic individuation, yet the format and flexibility of spatial scaffolding for working memory remain elusive. We hypothesised that information in working memory can be re-coded from its native format at encoding to organise and retain internal representations sparsely. To test this, we presented to-be-memorised visual objects at distinct directions and distances and leveraged gaze biases during mnemonic selection as an implicit read-out of spatial scaffolding for working memory. We report how the human brain abstracts away over incidental object distance when direction alone suffices as a scaffold, but incorporates distance when it aids mnemonic individuation. This unveils the principle of sparse spatial scaffolding for working memory, whereby the human brain flexibly resorts to the minimal spatial scaffold needed for the individuation of internal representations.

## Introduction

A central challenge for working memory is to code for information in a format in which individual representations remain separated and can be selectively prioritised for guiding behaviour [1–6]. Within the study of visual working memory, ample studies have made clear how space serves as a foundational scaffold for the separation and selection of individual working-memory representations [7–16]. Yet, how precisely space is used for working memory remains elusive and has been the subject of ample recent investigation.

Several recent studies have made clear how, once visual information has been encoded into working memory, the use of spatial location as a scaffold for memory individuation and selection is not necessarily veridical, but can be flexibly updated. For example, spatial organisation in visual working memory can be transformed [17–23], compressed [24], and engage additional spatial frames of reference [19,25,26]. This flexible nature of working memory – in which information can be re-coded from its native format at encoding (see also [23,27–33]) – provides the opportunity to tune the spatial scaffolding for working memory to the task in an adaptive and efficient manner. Accordingly, working memory may organise and retain mnemonic visual contents sparsely: utilising more sparse (abstracted) spatial scaffolds as long as they suffice for mnemonic individuation and selection, and engaging richer (more veridical) spatial scaffolds only when necessary.

To test this hypothesis of sparse spatial scaffolding for visual working memory, we developed a task in which we never tested the location of specific memory items, but in which space served as a medium for memory organization – or scaffold – serving item separation and selection. Critically, we presented visual memory items at different directions and distances from fixation, such that item distance was either useful as a spatial scaffold for mnemonic individuation (because direction alone was insufficient), or redundant as a spatial scaffolding feature (because direction alone was sufficient for mnemonic individuation).

To study spatial coding for visual working memory without asking participants about memorised item locations, we turned to spatial biases in fixational gaze behaviour as observed during mnemonic selection. This approach builds on prior research that has shown how spatial biases in fixational gaze behaviour can index spatial shifts of attention [34–38] even when shifting attention among visual contents in working memory [19,25,39–42]. Here we uniquely leveraged these peripherally observable spatially-indexed biases to probe the spatial scaffolding used for working memory implicitly, i.e., without ever asking participants about memorised locations. Doing so, we reveal the principle of sparse spatial scaffolding for working memory: exclusively utilising memorised item direction – and abstracting away over distance – when direction is sufficient for mnemonic individuation, and additionally incorporating item distance only when direction is insufficient for mnemonic individuation.

## Results

Human participants performed working-memory tasks in which we cued (via a central colour retro-cue) the selection of one of four spatially separated visual items for an orientation comparison to the ensuing test stimulus (Fig. 1a). We presented items either near (3 degrees) or far (6 degrees) from the central fixation dot and used spatial biases in fixational gaze behaviour during mnemonic selection as an implicit read-out of the spatial scaffolding used for visual working memory (an approach that we applied successfully in several recent complementary studies: [19,25,39,40,42]).

**Figure 1.**
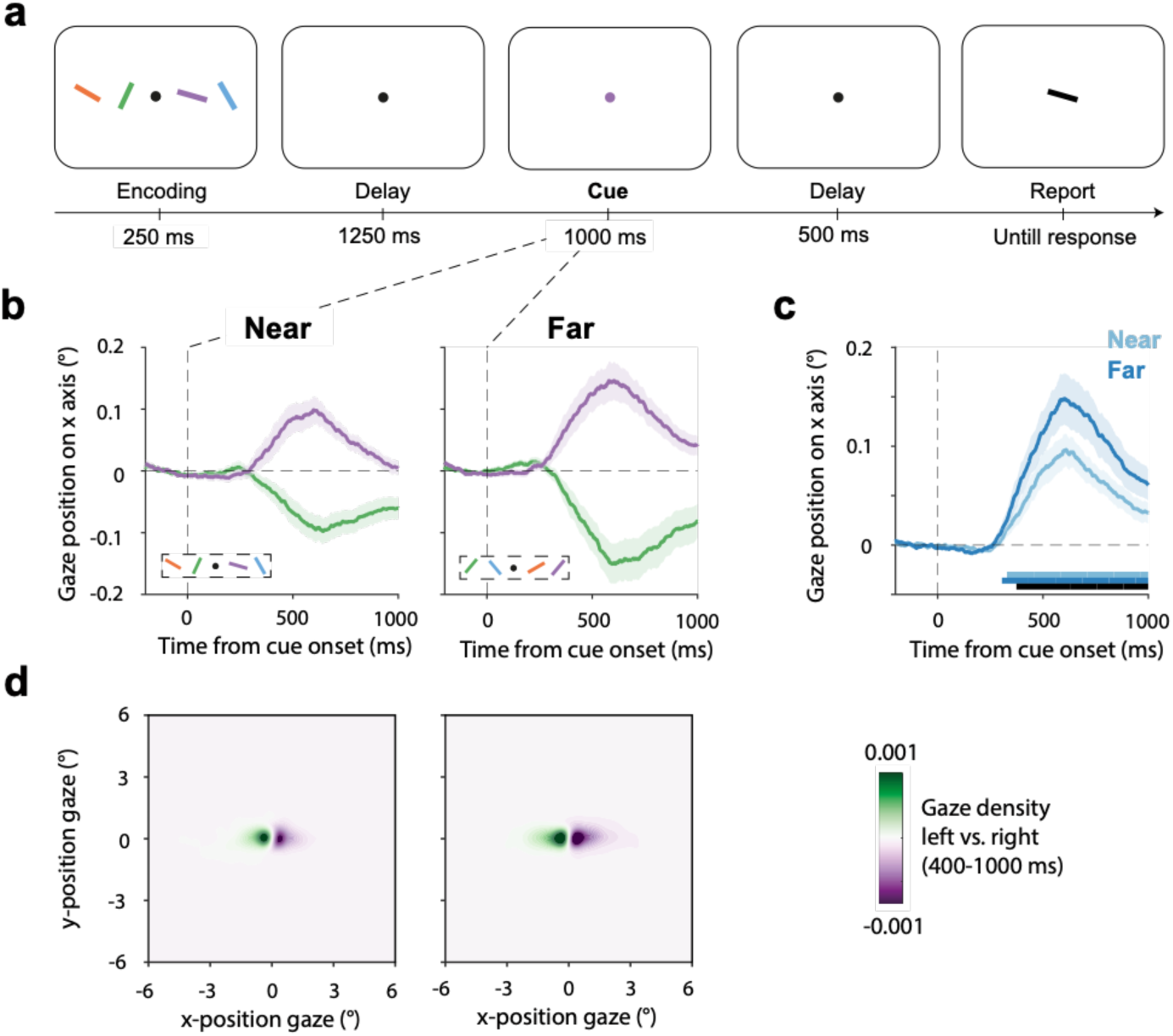
Fixational gaze behaviour during mnemonic selection tracks the use of both direction and distance as spatial scaffolding features for visual working memory. a) Task schematic. Participants memorised four bars with different colours and orientations. Following a delay, a colour change of the central fixation dot (retro-cue) prompted participants to select the colour-matching item from working memory to compare its orientation to a black bar in the upcoming test display. Memory items were presented to the left or right and near (3°) or far (6°) from fixation at encoding, but participants were never asked about memorised item location. b) Time course of average gaze position following central retro-cues that prompted the selection of the memory item that had been presented to the left or right at encoding, separately for when the memory item was near (left panel) or far (right panel) at encoding. c) Time courses of spatial biases in gaze toward the cued memory item when this item was near (light) or far (dark) at encoding. d) Heatmaps of two-dimensional gaze position following left vs. right cues. Shading in panels b-c indicate ±1 SEM calculated across participants (n = 25).

In what follows, we first establish that our spatial marker can track the use of both ‘direction’ and ‘distance’, when direction and distance are both useful as a spatial scaffold for individuating the four visual memory items – even when direction and distance were never asked about. We then show across two additional experiments that participants resort to a sparse spatial scaffold that abstracts away over distance, when direction alone is sufficient for individuating the four memory items.

The key outcome variable in our study was gaze during mnemonic selection. Before turning to our gaze data, we first confirmed that participants were able to perform this task well above chance in all three versions that we ran (Supplementary Fig. 1).

### Fixational gaze behaviour during mnemonic selection tracks the use of both direction and distance as spatial scaffolding features for visual working memory

Figure 1b shows gaze time courses after retro-cues that prompted the selection of memory items that had been encoded left or right from fixation, at either the near (Fig. 1b, left) or far (Fig. 1b, right) position. Consistent with prior studies from the lab, we observed clear spatial biases in gaze in the direction of the original location of the cued memory item. Importantly, this occurred even though (1) there was nothing to look at to the left and right after the retro-cue, (2) our colour retro-cue and ensuing test stimulus were presented centrally, and (3) we never asked participants about the original location of memory items .

To study whether distance was incorporated in the spatial scaffold used for working memory – in addition to direction – we collapsed left and right trials into a single measure of ‘towardness’ (as in [39,40,42,43]) and overlayed towardness between trials with cues to near and far items. As shown in Figure 1c, we found that gaze after the retro-cue became significantly biased in the direction of the cued memory item, both when the cued item was near (cluster P < 0.001) or far (cluster P < 0.001). In addition, the spatial bias in gaze was clearly modulated by the memory item’s distance at encoding, with a larger bias when cued to select the further item (cluster P < 0.001; black horizontal line in Fig. 1c).

The observed gaze biases originated from biases in fixational gaze behaviour (as in [34,35,42,43]) – and not looking back at the original location of the memorised item (as in [44–48]) at 3 or 6 degrees, respectively. This can be appreciated by the heat maps (two-dimensional density plots) of the difference in gaze position following cues to select the left vs. right items, as depicted in Figure 1d (see Supplementary Fig. 2 for the heatmaps of gaze density following left and right cues separately).

These spatial biases in fixational gaze behaviour during mnemonic selection confirm the use of space as a scaffold for mnemonic individuation, even if item locations are never asked about. These data further show that our gaze marker of this spatial scaffold is able, in principle, to track the use of both direction (left/right) and distance (near/far).

### Fixational gaze behaviour reveals a sparse spatial scaffold that abstracts over distance when direction alone is sufficient for mnemonic individuation

In Experiment 1, there were always two memory items in each direction (two left and two right). Accordingly, direction (left/right) alone was *insufficient* as a spatial scaffold for individuating the four items, rendering distance useful as an additional scaffolding feature. Following this logic, we refer to this condition as “direction-insufficient → distance-useful” (Fig. 2a, left).

**Figure 2.**
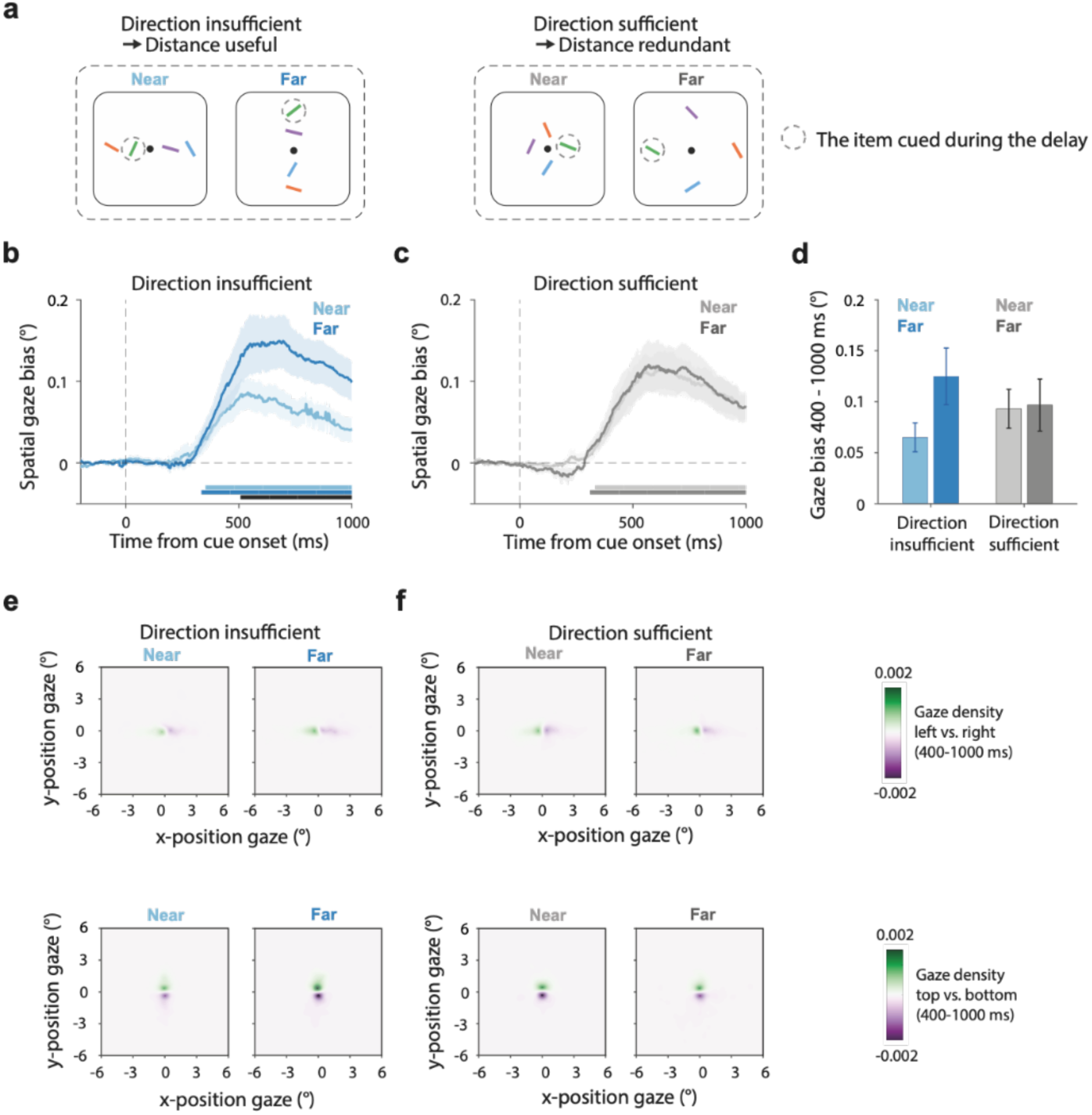
Fixational gaze behaviour reveals a sparse spatial scaffold that abstracts over distance when direction alone is sufficient for mnemonic individuation. a) Representative encoding displays used in Experiment 2. In Experiment 2 we additionally manipulated the usefulness of distance as a scaffolding feature for mnemonic individuation. The “direction-insufficient → distance-useful” condition (left) mirrored Experiment 1 where direction alone was insufficient for mnemonic individuating, rendering distance useful as an additional scaffolding feature. In the alternative “direction-sufficient → distance-redundant” condition (right), the four items each had a unique direction, rendering direction sufficient for item individuation and distance redundant as an additional scaffolding feature. Dashed circles indicate the later cued item in the different conditions (dashed circles were never shown in the experiment). b-c) Time courses of spatial biases in gaze toward the cued memory item when this item was near (light) or far (dark) at encoding, when direction is insufficient and distance useful as a spatial scaffold (panel b) or when direction was sufficient and distance redundant as an additional spatial scaffolding feature (panel c). d) Gaze bias averaged across the a-priori defined window from 400 to 1000 ms after the cue (based on [42]). e-f) Heatmaps of two-dimensional gaze position following left vs. right cues (top row) or top vs. bottom cues (bottom row) across our four conditions. Shading in panels b-c and error bars in panel d indicate ±1 SEM calculated across participants (n = 25).

In Experiment 2, we included an additional condition in which each memory item was placed along a separate direction from fixation (up, down, left, and right), while again manipulating item distance. Critically, in this condition direction became sufficient (in principle) as a spatial scaffold for individuating the four memory items, potentially rendering distance redundant as an additional scaffolding feature. Accordingly, we refer to this condition as “direction-sufficient → distance-redundant” (Fig. 2a, right).

When considering the condition in which distance was useful as an additional scaffolding feature because direction alone was insufficient (Fig. 2b), we replicated our findings from Experiment 1. We again found that gaze after the retro-cue became significantly biased in the direction of the cued memory item, both when the cued item was near (cluster P < 0.001) or far (cluster P < 0.001), and found that the spatial gaze bias was again clearly modulated by the memory item’s distance at encoding (Fig. 2b), with a larger bias when cued to select the further item (cluster P < 0.001).

Our key insight comes from the same spatial gaze bias in the condition in which all four memory items each had a unique direction (left, right, top, bottom). Here, we still observed clear gaze biases in the direction of the cued memory item (Fig. 2c), both when the cued item was near (cluster P < 0.001) or far (cluster P < 0.001). Critically, however, in this condition, we no longer found a modulation by distance (Fig. 2c). Importantly, this was observed even though we had presented the items at the exact same distances (3 and 6 degrees) as in the alternative condition. This lack of modulation by distance exclusively in this condition is consistent with the notion that, in this condition, direction was sufficient as a scaffold, affording the brain to “abstract away” over distance.

To further quantify this key finding, we collapsed these gaze biases over the pre-defined window of 400-1000 ms after cue onset – a window chosen a-priori based on [42]. This aggregate gaze-bias measure is depicted in Figure 2d. Analysis of variance confirmed a statistically significant interaction between the factors “whether the cued item was near or far” and “whether direction was insufficient (distance useful) or sufficient (distance redundant)” (F(1, 24) = 9.13, p = 0.006, partial η^2^ = 0.276). Post-hoc analysis confirmed that when direction was insufficient as a spatial scaffold for individuating all four memory items, participants had a larger gaze bias when selecting the far compared to the near memory item (t(24) =-2.872, p _Bonferroni_ = 0.05, Cohen’s d = −0.574). However, when direction was sufficient for individuating the four items (because each memory item appeared in a distinct direction from fixation at encoding), participants showed highly similar gaze bias regardless of whether we cued the near or the far memory item (t(24) =-0.401, p _Bonferroni_ = 1, Cohen’s d = 0.08). This suggests the use of a “sparser” spatial scaffold in this condition, in which item distance was redundant as an additional scaffolding feature.

Gaze heat maps, here split for trials with horizontal and vertical configurations, again revealed the fixational nature of these gaze biases (Fig. 2e-f).

### Our findings are driven by spatial scaffolding demands, not the mere presence of near and far items

In Experiment 2, we manipulated whether direction was sufficient (distance redundant) or insufficient (distance useful) as a spatial scaffold for individuating the four memory items by presenting items either in unique directions from fixation, or by presenting multiple items in the same direction from fixation. However, in the former condition, the four items were always *all* near or *all* far, while in the latter condition displays always contained two near and two far items. Accordingly, it remains possible that our findings of a distinct spatial scaffold between the two conditions reflects the mere presence of both near *and* far items in one condition, and only near *or* far items in the other condition.

To rule out this possibility, we designed Experiment 3 where we repeated our key manipulation, but this time ensured that displays always contained two near *and* two far items in both conditions (Fig. 3a). The direction-insufficient (distance-useful) condition mirrored Experiment 2 (Fig. 3a, left), while the direction-sufficient (distance-redundant) condition now always had two items placed at the ‘near’ positions on one axis and another two items placed at the ‘far’ positions on the orthogonal axis (Fig. 3a, right). Accordingly, both conditions now contained items at both distances, while direction was still sufficient (and distance redundant) as a spatial scaffold in one condition but not in the other.

**Figure 3.**
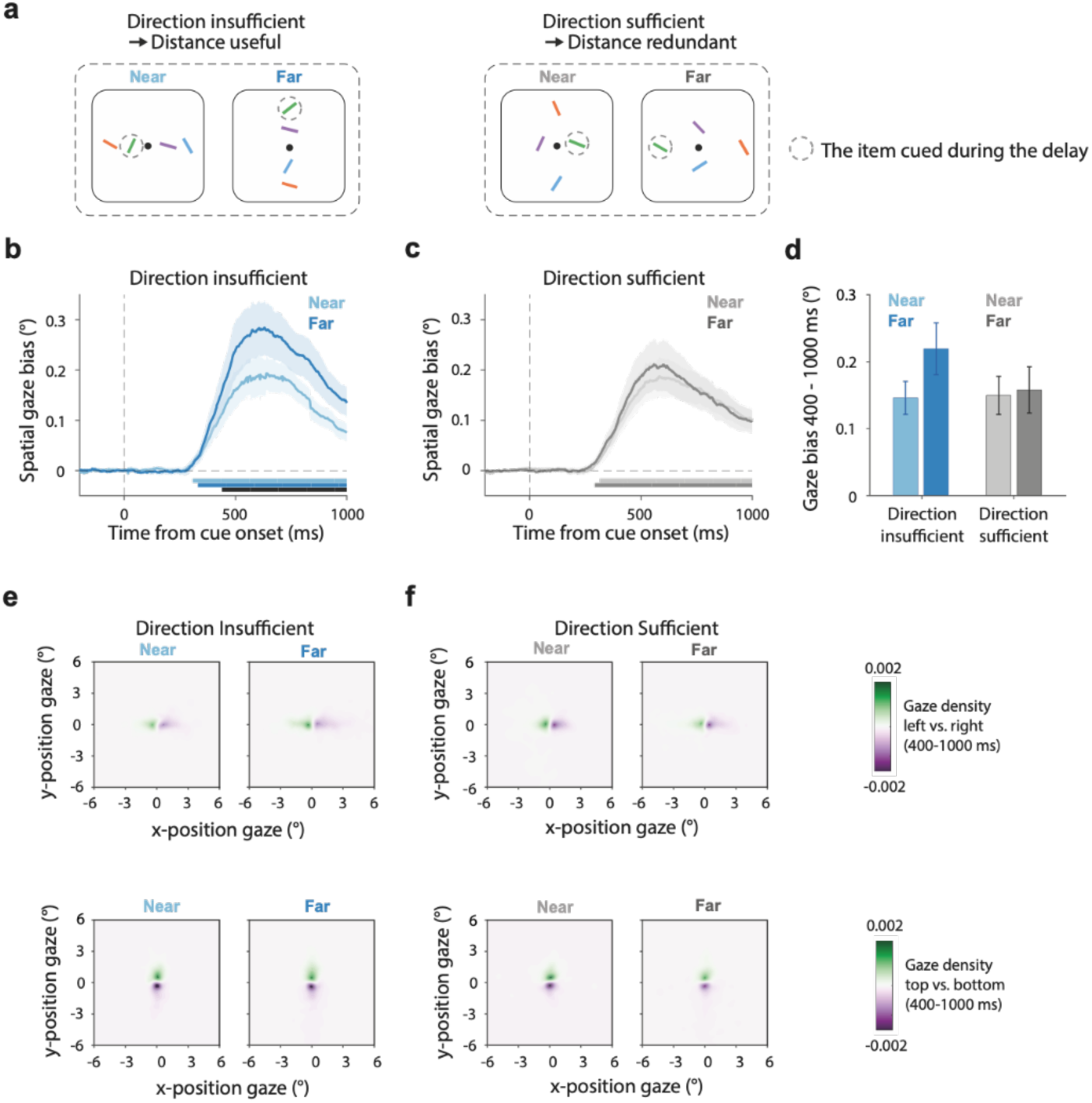
Our findings are driven by spatial scaffolding demands, not the mere presence of near and far items. a) Representative encoding displays used in Experiment 3. Experiment 3 replicated Experiment 2 except that in Experiment 3, the “direction-sufficient → distance-redundant” condition (right-top panel) now always also had two near items (on one axis) and two far (on the other axis). b-c) Time courses of spatial biases in gaze toward the cued memory item when this item was near (light) or far (dark) at encoding, when direction was insufficient and distance useful as a spatial scaffold (panel b) or when direction was sufficient and distance redundant as an additional spatial scaffolding feature (panel c). d) Gaze bias averaged across the a-priori defined window from 400 to 1000 ms after the cue (based on [42]). e-f) Heatmaps of two-dimensional gaze position following left vs. right cues (top row) or top vs. bottom cues (bottom row) across our four conditions. Shading in panels b-c and error bars in panel d indicate ±1 SEM calculated across participants (n = 25).

Gaze data in Experiment 3 replicated those from Experiment 2: again, showing a larger fixational gaze bias when selecting the further item, but only in the condition where direction was insufficient – and hence distance useful – for individuating the four memory items (Fig. 3b; cluster P < 0.001) and not in the direction-sufficient condition in which distance was redundant (Fig. 3c; no far-vs-near cluster). These findings were again corroborated by a significant interaction (Fig. 3d; F(1, 24) = 7.54, p = 0.011, partial η2 = 0.239), with post-hoc analysis confirming larger fixational gaze bias when selecting the further item when direction was insufficient (distance useful) for item individuation (t(24) =-3.518, p _Bonferroni_ = 0.011, Cohen’s d = −0.704), but not when direction was sufficient and distance was redundant as a scaffolding feature (t(24) =-0.55, p _Bonferroni_ = 1, Cohen’s d = −0.11). The heat maps in trials with horizontal and vertical configurations again revealed the fixational nature of these gaze biases (Fig. 3e-f).

## Discussion

We unveil how the human brain engages fundamentally distinct spatial codes for retaining visual representations in working memory, depending on the utility of spatial features as a scaffold for memory. Leveraging spatial biases in fixational gaze behaviour during mnemonic selection as an implicit read-out of spatial scaffolding for visual working memory (as in [19,25,39,40]), our present data unveil the principle of “sparse spatial scaffolding” for visual working memory, whereby the brain uses the minimal spatial features necessary for the separation and selection of individual contents in visual working memory.

Rather than investigating how we memorise space (cf. [26,29,49–53]), we asked how we *use* space for memorising. We never asked participants to report memorised item direction nor distance. Instead, space served as an organising medium (scaffold) serving the separation and selection of individual memory contents. While ample studies have made clear *that* space is a useful scaffold for working memory [7–14,54], *how* space is used has remained elusive. Inspired by several findings [27–29], we hypothesised that the brain may abstract away from the veridical spatial lay-lout at encoding to a more efficient or “sparse” spatial scaffold – as long as it serves the job of mnemonic individuation. We found evidence for this. In our set-up, this was achieved by abstracting away over distance when direction was sufficient for mnemonic individuation, but incorporating distance as an additional scaffolding feature when direction was insufficient (when multiple items existed along the same direction).

Ultimately, the strength of our inference hinges on the suitability of our spatial marker for studying spatial scaffolding for working memory. Our findings build directly on several prior studies where we successfully leveraged spatial biases in fixational gaze behaviour to study spatial scaffolding for working memory – without ever asking participants about memorised item location [15,19,25,39,40,42]. Here, we for the first time use this implicit marker to study the use of distance. In all three experiments, we confirm that this marker – while fixational in nature – is sensitive to memorised item distance, in principle. This suggests that the lack of a “distance code” when distance is a redundant scaffolding feature – our key finding – is not due to a lack of sensitivity of our marker, but rather reflects a sparser spatial scaffold in which distance has been abstracted away.

At the same time, the observed sensitivity to distance (when useful) reveals how biases in fixational gaze behaviour signal more than a low-level “orienting response” that merely reflects in which direction to orient – as may be predicted considering this gaze bias may reflect a spillover from activity in subcortical brain circuitry with evolutionary roots in orienting (cf. [55–57]). While remaining fixational in nature, our data reveal how this gaze bias is sensitive to “high-level” cognitive coding by tracking the flexible utilisation of distinct spatial scaffolds for working memory – that can either incorporate or abstract away over distance. This adds to growing appreciation of higher-level cognitive processing within deep-brain structures, such as the superior colliculus [58–61] that has been implicated in spatial biases in fixational gaze behaviour that we report here [62].

By studying spatial coding for working memory following a cue that directed internal selective attention to a specific memory item, our findings complement prior studies on selective attention that also varied item distance [34,63–70], even if these typically studied externally directed attention. Interestingly, at least several studies reported a relative invariance of spatial modulations in neural activity (e.g., [63–65]) or fixational gaze behaviour [34] to the attended target’s distance, while yet other studies did report distance-dependent modulations (e.g., [66–69]). We unveil a key variable that may underpin whether or not this is found: the utility of distance for the task at hand (see also [70]).

Though our starting point was different, our experimental manipulations resemble those in complementary research on the role of eccentricity and crowding in vision (e.g., [71–79]) and visual working memory (e.g., [80–82]). This literature has typically focused on other questions, such as how visual resolution changes when moving from central to peripheral vision. Our task was not designed to tap into such differences. We used clearly visible stimuli and placed them 3 degrees apart. Also, when considering performance, we did not observe lower accuracy for the far item in any of the three experiments (Supplementary Fig. 1). Instead, we here focused on whether and how distance was used as a scaffold for visual working memory when distance is a useful or a redundant scaffolding feature (in addition to direction). While our direction-insufficient (distance-useful) condition may invoke more crowding, we note that our key findings do not consist of a general increase or decrease of the fixational gaze bias in this condition (but rather an interaction with distance). Furthermore, the main insight from our study – that distance is abstracted away when it is a redundant scaffolding feature – originated from our condition with minimal crowding, when all items were presented in a distinct direction.

Having unveiled the principle of sparse spatial scaffolding for visual working memory, our data open relevant avenues for future research. Our data leave unaddressed how the spatial “pruning” of visual working memory develops between encoding and our moment of exposing the spatial scaffold after the cue, and how such spatial pruning occurs across the multitude of brain areas involved in working memory [83]. Our findings hint most directly at the spatial scaffold used by oculomotor circuitry in the brain, of which our gaze marker is a peripheral fingerprint [62]. Developing ways to continuously track the transformations in the spatial scaffold for working memory across time and across the brain remain challenging but also exciting avenues awaiting future research.

## Methods

### Ethics

Experimental procedures were reviewed and approved by the local Ethics Committee of the Vrije Universiteit Amsterdam. Each participant provided written consent before participation and was reimbursed €10/hour.

### Participants

We conducted three experiments with independent participant recruitment. Twenty-five healthy human volunteers participated in each experiment (Experiment 1: age range: 18-31; 6 male and 19 female; 23 right-handed and 2 left-handed; Experiment 2: age range: 18-27; 3 male, 21 female, and 1 non-binary; 24 right-handed and 1 left-handed; Experiment 3: 20-30; 7 male and 18 female; 22 right-handed and 3 left-handed). Sample size of 25 per experiment was set a-priori based on previous publications from our lab that relied on the same outcome measure [25,42,43]. In Experiment 3, to achieve the intended sample size, two participants were replaced due to the poor quality of the eye-tracking data. No participants required replacement in Experiments 1 and 2. All participants had normal or corrected-to-normal vision.

### Stimuli and procedure

We designed a visual working-memory task in which we never tested the location of specific memory items, but in which space served as a scaffold for item separation and selection. In the critical versions of our tasks, we presented visual memory items at different directions (left/right/top/bottom) and distances (3 or 6 degrees) from fixation, and manipulated the spatial lay-out such that item distance was either useful as a spatial scaffolding feature (because direction alone was insufficient) or redundant (because direction alone was sufficient). Below, we first explain the basic task, before returning to our two key manipulations.

In all three experiments, participants engaged in a visual working-memory task that required the selection of a visual item from working memory for an upcoming orientation comparison (Fig. 1a). Though we manipulated spatial features of the display, note how we never asked participants about memorised item direction, nor about memorised distance. To perform our task, memories of colour-orientation bindings were sufficient, in principle. Spatial location merely served as a scaffold for separating and selecting specific memory items.

Each trial began with a start fixation (200 ms) followed by a brief (250 ms) encoding display where four bars with different colours and orientations appeared on either two or four directions of the fixation dot. After a 750 ms working-memory retention interval, the fixation dot changed colour for 1000 ms serving as a retro-cue. This retro-cue cued with 100% validity which memory item (i.e. the colour-matching item) would have to be compared to the upcoming test stimulus. The retro-cue was followed by another retention delay of 500 ms before the test display appeared. During the test display, a black bar appeared at the center of the screen. The test bar was always rotated between 10 to 20 degrees clockwise or counter-clockwise from the cued bar in memory. Participants reported whether the cued memory item should be turned clockwise or counter-clockwise to match the black bar in the test display. Participants received feedback immediately after response by a number (“0” for wrong, or “1” for correct) appearing for 250 ms slightly above the fixation dot. After the feedback, inter-trial intervals were randomly drawn between 500 and 1000 ms.

Our key experiments (Experiments 2 and 3) had two key manipulations (Fig. 2a; Fig. 3a). We manipulated: (1) whether cued memory items were presented near (3°) or far (6°) from fixation, and (2) whether all items were presented in a unique direction (left/right/bottom/top) or whether certain directions were shared between multiple items in the array (two left and two right items, or two top and two bottom items). Critically, in the first condition (Fig. 2a, labelled “direction-insufficient → distance-useful”), there were always two items competing along each direction of an axis (e.g., two up and two down, or two left and two right), rendering direction alone insufficient as a spatial scaffold for individuating the four items, and rendering distance useful as an additional scaffold. In contrast, in the alternative condition (Fig. 2a, labelled “direction-sufficient → distance-redundant”), all items were associated with a unique direction, such that direction was sufficient as a spatial scaffold for individuating the four memory items, rendering distance redundant..

Experiment 1 (Fig. 1a) only included the condition with four items on a line (direction-insufficient → distance-useful) and served to validate the sensitivity of our spatial marker to track the use of both ‘direction’ and ‘distance’ when both features were useful as a scaffold. In Experiments 2 and 3, we added the direction-sufficient (distance-redundant) condition in which all items were presented in a different direction from fixation (see Fig. 2a and Fig. 3a). In Experiment 2 (Fig. 2a), in this direction-sufficient (distance-redundant) condition, the display in a given trial always contained four items that were either all near (3 degrees) or all far (6 degrees). In contrast, in Experiment 3 (Fig. 3a) displays in this condition always contained two near items (one axis) and two far items (on the orthogonal axis), to equate the total number of near and far items in each display between the two conditions in which direction was either sufficient (and distance redundant) or insufficient (and distance useful) as a spatial scaffold for item individuation. Please note that in Experiment 1, we only included the horizontal configuration (as depicted in Fig. 1a), while in Experiments 2 and 3, we included both vertical and horizontal configurations in the direction-insufficient (distance-useful) condition (as depicted in Fig. 2a; Fig. 3a), such that we equally often cued a left/right/top/bottom item in both the direction-insufficient (distance-useful) and direction-sufficient (distance-redundant) conditions.

In the encoding display, bars were randomly assigned two or four distinct colours from the colour pool: green (RGB: 133, 194, 18), purple (RGB: 197, 21, 234), orange (RGB: 234, 74, 21), and blue (RGB: 21, 165, 234]). Bars were also drawn at distinct orientations ranging from 0° to 180° with a minimum difference of 20° between each other. During the test display, the bar were always dark grey (RGB: 64, 64, 64) and were always oriented clockwise or counter-clockwise from the cued memory target, with a change in orientation randomly drawn between 10 to 20 degrees. The bar is 2 visual degrees in length and 0.4 visual degrees in width and the fixation point has a radius of 0.07 visual degrees.

Experiment 1 consisted of 4 sessions that each contained 10 blocks of 16 trials, resulting in a total of 640 trials. In Experiments 2 and 3, we added the direction-sufficient (distance-redundant) condition and therefore increased the number of trials. Experiments 2 and 3 each consisted of 5 sessions, which each contained 5 blocks of 32 trials, resulting in a total of 800 trials. Conditions were randomly mixed within each block.

Before the start of each experiment, participants practiced the task for approximately five minutes. Experiment 1 required approximately 80 minutes while Experiments 2 and 3 each required approximately 100 minutes per participant to complete. Participants were instructed to keep fixation throughout the task, but trials were not aborted when larger eye movements occurred. The encoding display was too brief to allow gaze fixations to all memory items, and during the main cue-period of interest there was nothing to look at on the screen apart from the central fixation dot.

### Eye tracking

We used an EyeLink 1000 (SR Research, with 1000 Hz sampling rate) to track gaze from a single eye (right eye in all participants except 1 for which the left eye provided a better signal). The eye-tracker camara was positioned on the table approximately 5 cm in front of the monitor and approximately 65 cm in front of the eyes.

Gaze data were read into Matlab using the Fieldtrip toolbox [84]. In line with previously established protocols [42,43,85], we first identified blinks and replaced identified blink clusters (extending 100 ms before and after detected blinks) with Not-a-Number (NaN) to effectively mitigate any artifacts from the blinks. Then, data were epoched from-1000 to +2000 ms relative to the onset of the retro-cue.

We focused our analysis on spatial biases in gaze position over time in response to the central colour retro-cue that prompted the selection of one out of the four items in working memory. Because the cue was non-spatial and the test stimulus would occur centrally, any spatial bias in gaze during this period must reflect selection within the spatial lay-out of visual working memory.

To investigate directional biasing of gaze during mnemonic selection, we first baseline corrected the eye data by subtracting the mean data in the [-200 to 0 ms] baseline window. Then, we examined eye data on the x-axis when the cued item was on the left or the right side, on the y-axis when the cued item was above or below fixation. Following our prior studies [25,42,43], we condensed the relevant data from the left/right and top/bottom trials into a single gaze time course of “towardness” that captured the bias of gaze towards the memorized location of the cued memory item. We could then compare this measure of towardness across our experimental conditions.

To zoom in on biases in fixational gaze behaviour, we removed occasional trials with large gaze deviations from fixation (as was also done in [40,43,86]) within the period of interest from 0-1000 ms relative to cue onset. Specifically, we removed trials in which gaze values were larger than 2 degrees away from fixation, which corresponded to the inner radius of the near item positions. This ensured that all reported effects cannot be driven by looking at the original location of the cued memory items, but instead are fixational in nature. Gaze remained close to fixation in the majority of trials, hence relatively little trials were removed following this procedure: (usable trials: Experiment 1: 95 ± 1.4%, Experiment 2: 93 ± 1.7%, Experiment 3: 91 ± 2.6%).

In addition, the fixational nature of the gaze bias was also visualized through 2-dimensional (2D) heat maps of gaze density (as in [25,42,43]). For this, we collated gaze-position values across time and trials (without averaging) using the data from the 400 to 1000 ms window after the retro-cue (a time window set a-priori based on [42]). We then counted the number of gaze samples within 0.1° x 0.1° bins, ranging from-6° to 6° and sampling the full 2D space in steps of 0.05°× 0.05°. To obtain a density map, we divided 2D gaze-position counts by the total number of gaze-position samples. For visualization purpose, density maps were smoothed using a 2D Gaussian kernel with an SD of 0.25° × 0.25° (using the built-in function “imgaussfilt” in MATLAB). To provide and comprehensive and undistorted view of gaze density in our task, trials with large gaze deviations were not removed from the data entering these gaze-density visualisations.

To represent the heat map associated with the directional gaze biases of interest, we first obtained the maps separately following cues to left, right, top, and bottom items and subtracted maps between conditions in which cues were associated with items in the opposite direction: left versus right and top versus bottom. We did this separately for each of our core experimental conditions.

### Statistical analysis

To evaluate and compare gaze towardness time courses, we employed a cluster-based permutation approach [87], which is well-suited for evaluating multiple neighbouring time points while avoiding multiple comparisons. In this approach, we generated a permutation distribution by randomly permuting trial-average time courses at the participant level 10,000 times and identifying the largest clusters after each permutation. P-values were computed as the proportion of permutations where the largest post-permutation cluster exceeded the size of the observed cluster(s) in the original (non-permuted) data. Using Fieldtrip, we performed this permutation analysis with default cluster settings: grouping similarly signed significant data points from a univariate t-test at a two-sided 0.05 alpha level, and defining cluster size as the sum of all t-values within the cluster.

Because our central question regarded spatial scaffolding for working memory, we focused on the spatial gaze bias as our primary outcome variable because this is a direct spatial index. For completeness, we also considered our behavioural performance data as a function of our conditions. We note however that performance data in the current study mainly served to ensure that participants were able to complete the task across our conditions. We had no specific hypotheses about these data. Behavioral-performance data (accuracy and reaction times) were statistically evaluated using a two-way repeated-measures ANOVA with the factors whether the cued item was near or far and whether direction was sufficient (and distance-redundant) or direction was insufficient (and distance-useful) as a scaffold for separating the four items in working memory. ANOVA results were supplemented with post-hoc t-tests. As measures of effect size, we used partial eta squared for ANOVA and Cohen’s d for follow-up t-tests. P-values of follow-up t-tests were Bonferroni corrected for multiple comparisons.

## Data availability

All data will be made publicly available before publication.

## Code availability

Relevant code associated with the here-presented analyses will be made available through GitHub before publication.

## Acknowledgments

This research was supported by an ERC Starting Grant from the European Research Council (MEMTICIPATION, 850636) and an NWO Vidi Grant by the Dutch Research Council (grant number 14721) to F.v.E.

---

**Supplementary Figure 1.**
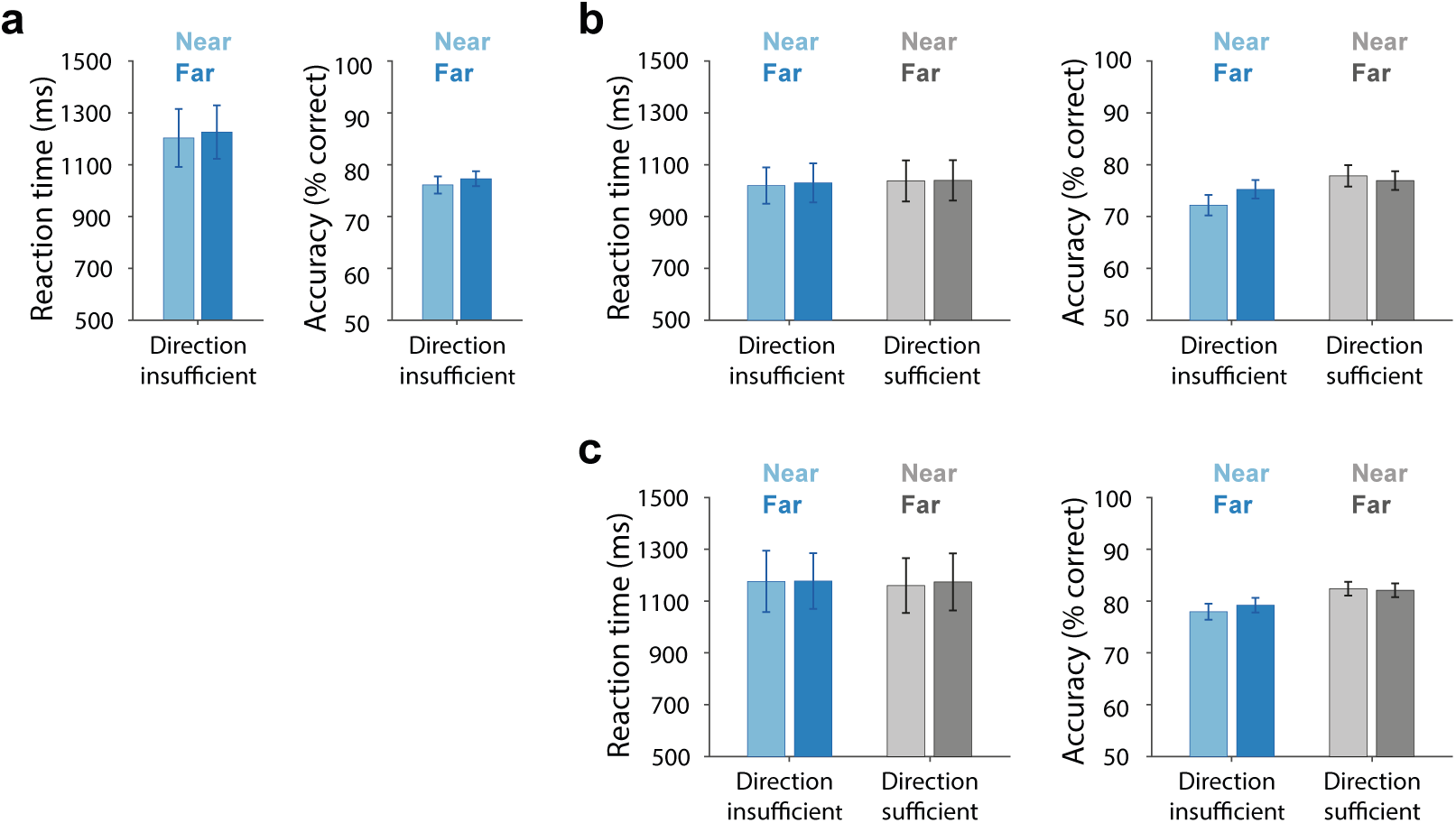
Largely comparable performance regardless of whether we cued the near or the far memory item in Experiments 1-3. a) In Experiment 1, participants showed similar performance regardless of whether we cued the near or the far memory items (Reaction time: t(1, 24) = −1.1, p = 0.28, Cohen’s d = −0.219; Accuracy: t(24) = −1.23, p = 0.23, Cohen’s d = −0.246). b) In Experiment 2, for reaction time, there is no significant main effect of direction-sufficiency (F(1, 24) = 0.86, p = 0.36, partial η2 = 0.035) nor on item distance (F(1, 24) = 0.33, p = 0.57, partial η2 = 0.013). We also did not observe an interaction (F(1, 24) = 0.08, p = 0.78, partial η2 = 0.003). For accuracy, we found a significant main effect of direction-sufficiency (F(1, 24) = 29.02, p < 0.001, partial η2 = 0.547), but no main effect of item distance (F(1, 24) = 3.16, p = 0.09, partial η2 = 0.116). Here, we did find a significant interaction (F(1, 24) = 9.13, p = 0.006, partial η2 = 0.276). Post hoc analysis suggests that when direction was insufficient as a spatial scaffold for item individuation, participants had higher (not lower) accuracy when selecting the far item compared to the near item (t(24) = −3.4, p Bonferroni = 0.014, Cohen’s d = −0.68). However, when direction was sufficient (and distance useful) as a scaffold, participants showed similar performance regardless of whether we cued the near or the far memory item (t(1, 24) = 1.05, p Bonferroni = 1, Cohen’s d = 0.21). c) In Experiment 3, for reaction time, there is no significant main effect of direction-sufficiency (F(1, 24) = 0.452, p = 0.508, partial η2 = 0.018) nor of item distance (F(1, 24) = 0.417, p = 0.525, partial η2 = 0.017). We also did not observe an interaction (F(1, 24) = 0.157, p = 0.695, partial η2 = 0.007). For accuracy, we again found a significant main effect of direction-sufficiency (F(1, 24) = 31.786, p < 0.001, partial η2 = 0.57), but no main effect of item distance (F(1, 24) = 0.476, p = 0.497, partial η2 = 0.019). Unlike in experiment 2, we did not find an interaction in Experiment 3 (F(1, 24) = 2.728, p = 0.112, partial η2 = 0.102).

**Supplementary Figure 2.**
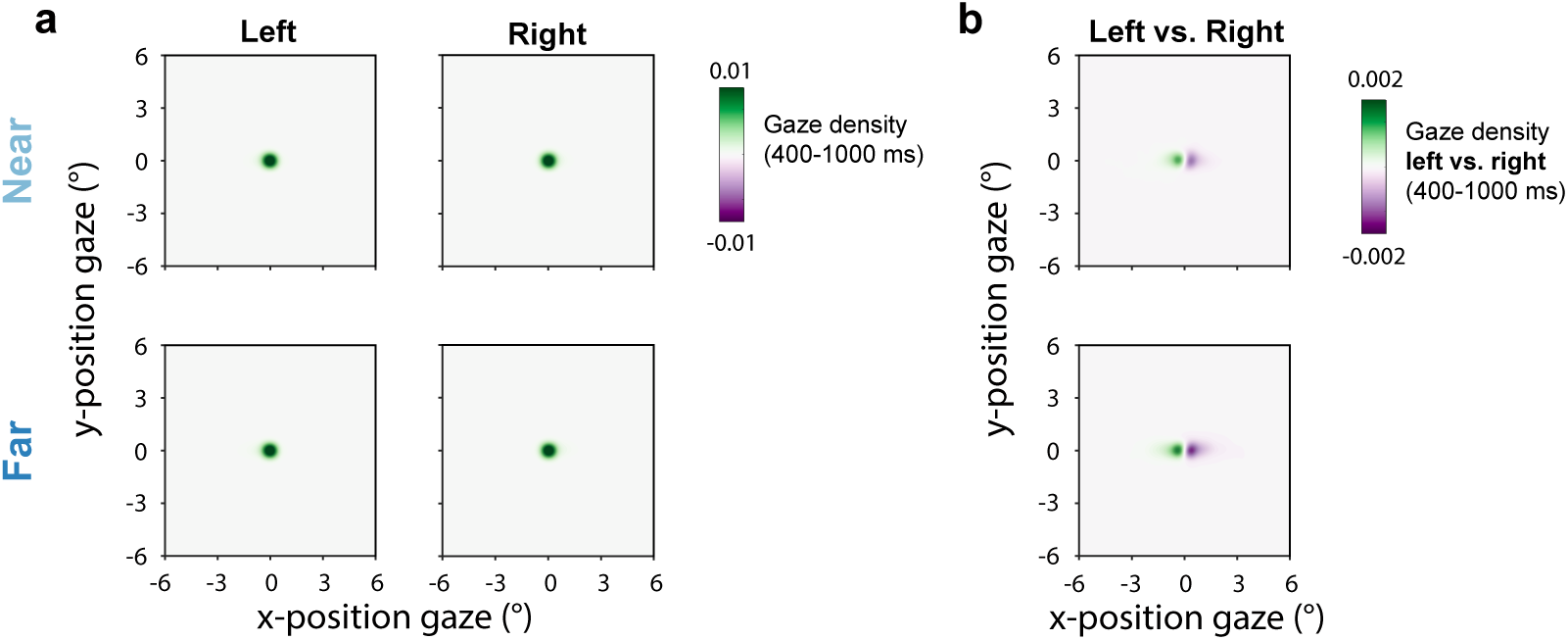
Complementary visualisation to the heat-maps of gaze density in main Figure 1d. a) Gaze density separately following cues that directed internal attention to the left and the right memory items. b) Difference in gaze density following cues directing attention to left vs. right items. Top row shows selection of ‘near’ items, while the bottom row shows selection of ‘far’ items. Data from Experiment 1.

